# Developing a broadly applicable *pyrF*-based genetic manipulation system in *Acinetobacter baumannii*

**DOI:** 10.1101/2020.11.05.370742

**Authors:** Run Xu, Can Gao, Shuqi Wu, Mengjiao Su, Chengfu Sun, Xu Jia, Rui Wang

## Abstract

*Acinetobacter baumannii* is an emergency pathogenic bacterium for its multidrug-resistance and high mortality rates after infection. In-depth genetic analysis of *A. baumannii* virulence and drug-resistant genes is highly desirable. Existing methods for genetic manipulation of *A. baumannii* mainly rely on the use of antibiotic as the selectable marker, and the *sacB*/sucrose as the counter-selectable marker, which is inconvenient and inappropriate for all research of *A. baumannii*. Based on the highly conserved *pyrF* gene and its conserved 500bp-flanking sequence, we quickly and easily generated the *pyrF*-deleted mutants as the uracil auxotrophic host strain in three model strains and 11 clinical strains. The *pyrF*-carried vectors constructed for gene editing with pyrF/5-FOA as the counter-selection were conveniently and time-saving in these *pyrF*-deleted mutants. Utilizing the *pyrF*-based genetic manipulation system, we easily and efficiently modified the *cas* gene and CRISPR sequence of I-F CRISPR-Cas system in *A. baumannii* AYE, and detected the CRISPR interference and adaptation in these mutants. In summary, the *pyrF*-based genetic manipulation system could be broadly applicable used for efficiently maker-less gene editing in most *A. baumannii* strains.

## Introduction

*Acinetobacter baumannii* (*A. baumannii*) is a most commonly and emergence pathogenic bacteria involved in hospital infection that causes wound infection, osteomyelitis, respiratory infections, bacteremia, etc. Since 1980s, multi-drug-resistant *A. baumannii* has become a new challenge in the treatment of nosocomial infections, thus the drug-resistance mechanism of *A. baumannii* has always been concerned [1]. Over the last decade, the resistant rate of *A. baumannii* has been increasing, and resistant strains of carbapenems have been widely reported [2–5]. Tigecycline and colistin have once been considered as the last antibiotics used to treat carbapenem-resistant *A.baumannii*[6, 7]. But unfortunately, tigecycline- and colistin-resistant *A. baumannii* has also appeared recently [8, 9]. The virulence mechanism of *A. baumannii* is also a focus of research in recent years as its high mortality rates observed for infected patients in the hospital environment[10]. Therefore, it is urgent to explore the molecular mechanism of drug-resistance, cytotoxicity and virulence of multidrug-resistant *A. baumannii*.

Uncovering the mechanisms of *A. baumannii* drug resistance and virulence required large amounts of genetic analysis. Once, the lack of genetic manipulation tools and methods for studying these issues has largely hindered the understanding of the diseases caused by *A.baumannii*. In the last several years, various putative methods for transform exogenous DNA into *A. baumannii* has been developed and optimized, like conjugation, electro transformation, and natural transformation via motility [11–14]. Several vectors were also been designed and modified for the genetic manipulation of *A. baumannii*, such as nonreplicative plasmids, replicative plasmids, or helper plasmids [12–18]. These vectors can be utilized to introduce DNA sequences directly into the chromosome of *A. baumannii* via single-crossover homologous recombination or complementation on a replicative plasmid with an antibiotic selectable marker. For multi-drug-resistant *A. baumannii* strains, it was usually not so easy to find a suitable antibiotic marker for a certain strain. However, tellurite as a non-antibiotic selectable marker was found to be broadly applicable used in most *A. baumannii* strains[15], and more non-antibiotic selectable markers also needed to be developed for research.

For constructing the clean and unmarked mutants, a counter-selectable marker was usually utilized to selectively screen the second-crossover colonies with the gene or suicide plasmid eliminated from the genome, or the replicative plasmid cured. Common to most methods in *A. baumannii* is that they used *sacB*/sucrose as a counter-selection system or natural lost the plasmids [14, 15, 17–19], which is broadly used in many organisms. The *sacB* gene expressed levansucrase in the presence of sucrose is lethal in some Gram-negative bacteria [20]. But when used the vectors with *sacB*/sucrose system in *A. baumannii* for screening the clean and unmarked mutations, it always needed to be subculture daily for several days with sucrose [15, 17]. This may due to some problem with the expression of *sacB* gene or the function of levansucrase in *A. baumannii*. So another counter-selection marker could be considered to be introduced into the vectors for efficiently gene editing in *A. baumannii*.

Auxotrophic makers, such as amino acid auxotrophy, NADPH auxotrophy, thymidine auxotrophy and uracil auxotrophy, are useful tools in gene editing and vaccine products [21–25]. The uracil auxotrophy has been successfully constructed in multiple strains, and some uracil related genes, such as *pyrF* and *ura3*, are also widely used [23, 26–28]. Uracil nucleotides are indispensable in the process of bacterial life, and orotate is a key intermediate in the synthesis of uracil nucleotides. The *pyrF* gene encodes orotate phosphate ribosyl transferase, which could catalyze the combination of orotate and phosphate ribose to produce orotate nucleotides. If the *pyrF* gene is deleted, the function of autonomously synthesizing uracil in bacterial life will be lost; bacteria could only use exogenous uracil nucleotides, and could not grow in medium lacking uracil nucleotides. 5-fluoroorotic acid (5-FOA), an analogue of orotic acid, can be converted to a highly toxic compound (5-fluoro-UMP, 5-F-UMP) by the product of *pyrF* [28]. Therefore, the *pyrF* gene can be used as the selection and counter-selection marker in the *pyrF*-deleted mutants.

Here, based on the pMo130Tel^R^ system[15] with tellurite and 5-FOA as the selectable and counter-selectable agent, we constructed the *pyrF*-deleted mutants as the uracil auxotrophic host strains in both *A. baumannii* model strains and clinical strains. Inaddition, the *pyrF* gene of *A. baumannii* AYE as the selectable and counter-selectable marker was introduced to generate the suicide plasmids for homologous recombination, and the replicative plasmids for gene expression in those strains. As results, the second-crossover colonies screened by *pyrF*/5-FOA system were more efficiently other than *sacB*/sucrose system, and the uracil auxotrophic medium M95/*pyrF* system also worked well as the selectable marker. Using this genetic system, we conveniently and efficiently modified the DNA sequences of I-F CRISPR-Cas system in *A. baumannii* AYE, and detected the interference and adaptation for these mutants. We found that all Cas proteins except Cas1 were involved in interference, and almost all Cas proteins and a priming spacer were needed for adaptation. So the *pyrF*-based genetic manipulation system could be conveniently and broadly constructed and used in most *A. baumannii* strains.

## Materials and methods

### Bacterial strains, plasmids and culture conditions

The strains and plasmids used in this study can be found in Table S1. *E. coli* strains were grown at 37°C in Luria-Bertani (LB) medium, supplemented with the appropriate agent. *A. baumannii* strains were grown at 37°C in LB medium, supplemented as needed with potassium tellurite (Tel, 30 mg/L), kanamycin (K, 50 mg/L), uracil (U, 50 mg/L), 5-Fluoroorotic Acid (5-FOA, 50 mg/L) and agar (15 g/L). For pyrimidine-free medium, a synthetic medium (M95) was used[28], which comprised 0.6% Na_2_HPO_4_, 0.3% KH_2_PO_4_, 0.1% NH_4_Cl, 1% NaCl, 1*10^−5^% thiamin-HCl, 0.2% glucose, 1 mM MgSO_4_, 0.5% Bacto casamino acids (pH 7.0).

### Molecular biology and bioinformatics methods

The gene sequences analyzed in this study were obtained from NCBI (https://www.ncbi.nlm.nih.gov/), amino acids and nucleotides sequences for *A. baumannii* were analyzed through BLASTP and BLAST (https://blast.ncbi.nlm.nih.gov/Blast.cgi). The primers used for vector construction and PCR detection are listed in Table S2. PCR was performed according to the manufacturer’s instructions by using the Phanta Max Master Mix (Vazyme) for cloning and 2×Taq Master Mix (Vazyme) for detection. Restriction enzymes used for cloning were from New England BioLabs (Ipswich, MA). Plasmids were all constructed through a homologous recombination technology using ClonExpress II One Step Cloning Kit (Vazyme).

### Plasmid construction

#### Construct plasmid for the *pyrF* gene deletion

The pMo130Tel^R^-DF vector was designed to delete the entire open reading frame of the *pyrF* gene in *A. baumannii*. The flanking upstream and downstream 500bp-regions of *pyrF* gene for *A. baumannii* AYE were PCR amplified and cloned into pMo130Tel^R^ (BamHI and PstI digested) to generate pMo130Tel^R^-DF.

#### Construct suicide plasmids with the *pyrF* gene as a selectable and counter-selectable marker

The pMo130TF and pMo130F vector were suicide plasmid for homologous recombination, which used the *pyrF* gene as the selectable and counter-selectable marker. The *pyrF* gene (along with its 200bp-upstream promoter sequence) of AYE and the *Km^R^* gene of pMo130 were PCR amplified and cloned into pMo130Tel^R^ (EcoRI and BamHI digested) or pMo130 (EcoRI and BamHI digested) to generate pMo130TF or pMo130F, in which the *sacB* gene was replaced by *pyrF* gene.

#### Construct autonomous replication plasmids with the *pyrF* gene as a selectable marker

The pMo130TFR or pMo130FR were autonomous replication plasmids, which did not need to integrate into genome and could use the *pyrF* gene as a selection marker. The *rep* gene of pWH1266[29] was PCR amplified and cloned into pMo130TF (EcoRI digested) or pMo130F (EcoRI digested) to generate pMo130TFR or pMo130FR.

#### Construct plasmids for gene manipulation

For gene deletion, the flanking upstream and downstream 500bp-regions of *cas1*, *cas3*, *cascade* and CRISPR were PCR amplified and cloned into pMo130TF (BamHI and PstI digested) or pMo130Tel^R^ (BamHI and PstI digested) to generate pMo130TF-E*cas1*, pMo130TF-E*cas3*, pMo130TF-E*cade*, pMo130TF-ECRI, pMo130Tel^R^-E*cas1*, pMo130Tel^R^-E*cas3*, pMo130Tel^R^-E*cade*, pMo130Tel^R^-ECRI. For gene insertion, the flanking upstream and downstream fragments of *cas3* termination codon and the GST tag of pGEX-6P-1 were PCR amplified, these three DNA fragments were then cloned into pMo130TF (BamHI and PstI digested) to generate pMo130TF-E*cas3-GST*.

#### Construct plasmids for CRISPR interference and adaptation detection

The pMo130TFR-ECC-sp2 was a targeted plasmid for detecting the CRISPR interference and adaptation. The PAM 5’-CC and spacer2 of AYE CRISPR were PCR amplified and cloned into pMo130TFR (NotI and BamHI digested) to generate the pMo130TFR-ECC-sp2.

#### Preparing the competent cells and electro transformation the plasmids

The preparation of *A. baumannii* competent cells for electro transformation was carried out as previously described [18], with some modifications. The *A. baumannii* to be electrically transformed was cultured to late-logarithmic phase in 2-3 mL liquid LB medium containing uracil. The cultures were centrifuged at 10,000 rpm and 4°C for 2 min. The harvested cells were washed three times with 10% cold glycerol, and then resuspended in 100μl of 10% cold glycerol for transformation.

For electrical transformation, the autonomously replicating plasmid pMo130TFR and its derivative plasmids need 1 μg to be mixed with the 100 μl competent cells, while the suicide plasmids and its derivative plasmids need 10 μg. The mixed cells were incubated on ice for 10 min, and then transferred to a 1 mm cuvette with Voltage 1800 V, Capacitance 25 μF, and Resistance 200 Ω, Cuvette 1mm. The competent cells after electroporation were transferred to a 1.5 mL EP tube with 600 μL LB liquid medium added, and cultured at 200 rpm and 37 °C for 40-60 min. The cells were plated on Tel-containing LB plates or M95 plates and cultured overnight at 37 °C.

### Isolation and characterization of the mutants

After the electrical transformation for plasmids, the cells were plated on Tel-containing LB or M95 plates to select the colonies which contained plasmids integrated into the genome or autonomous replicated. For gene deletion or insertion, the first-crossover colonies were then cultured overnight in liquid LB containing 50mg/L uracil and 50mg/L 5-FOA (or 10% sucrose), and subsequently diluted on the LB plates containing 50mg/L uracil and 50mg/L 5-FOA (or 10% sucrose). 20 colonies or more were picked to streak on the photocopying plate, for confirming the *pyrF* genes were eliminated from the genome. Subsequently, PCR was amplified to screen the colonies with correct size band.

### Detection of the CRISPR interference and adaptation

The pMo130TFR (control) and pMo130TFR-ECC-sp2 were electro transform into *A. baumannii* competent cells with the same condition. For CRISPR interference detection, we counted the colonies and conducted comparative analysis. For CRISPR adaptation detection, we used the primers to PCR amplify the sequence flanking the CRISPR leader side to detect the expanded band, relative to the parental band (control).

### Measuring the growth rates and sensibility to 5-FOA

The wild type strains and mutants of *A. baumannii* were streaked on both LB plates and LB plates with 5-FOA (50mg/L) added and culture overnight to test the sensibility to 5-FOA. For the uracil prototrophic of *A.baumannii*, we measured the growth curve by liquid cultured in both LB and LB with uracil (50mg/L) added, and also streaked on the M95 plates and M95 plates with uracil (50mg/L) added.

## Results

For construction of a *pyrF*-based genetic manipulation system, the *pyrF* gene was firstly deleted to generate *ΔpyrF* mutants as the uracil auxotrophic host strain for plasmids transform. And then for complement the function of *pyrF* gene in the *ΔpyrF* mutants, the *pyrF* gene should be expressed with the *pyrF*-carried plasmids transformation. Therefore, a suicide plasmid should be constructed with *pyrF* gene introduced, and could be used to conduct derivative plasmids for integrating into genome by homologous recombination; and a replicative plasmid also needed to be generated for autonomous replication without integrating into the chromosome. The former was usually used to delete, insert, and mutant the DNA sequence, while the latter was used for gene expression or complement.

### Identification of the *pyrF* gene and sensitivity to 5-FOA of *A. baumannii*

#### The *pyrF* gene and its flanking DNA sequences are highly conserved in *A.baumannii*

The *pyrF* gene sequence of *A. baumannii* was obtained from NCBI, and revealed that it encoded a 232-aminoacid polypeptide and was annotated as orotidine 50-phosphate decarboxylase. The *pyrF* gene sequences and its 500bp-flanking DNA sequences in most *A. baumannii* are highly conserved with the percent identity almost above 98% by NCBI blastn (https://blast.ncbi.nlm.nih.gov/Blast.cgi) (Fig. 1A). Based on this, a single plasmid contained the highly conserved 500bp-flanking DNA sequences would be generated to delete the *pyrF* gene in most *A. baumannii* strains, so as to broadly generate the uracil auxotrophic host. In addition, the *pyrF*-carried plasmids could be also constructed to be broadly used for genetic manipulation in these *ΔpyrF* mutants. Thus, in order to develop a broadly applicable *pyrF*-based genetic manipulation system in *A.baumannii*, three model strains *A. baumannii* str. AYE, ATCC17978, ATCC19606, and 11 clinical strains were selected for study (listed in Table S1).

**Figure 1.**
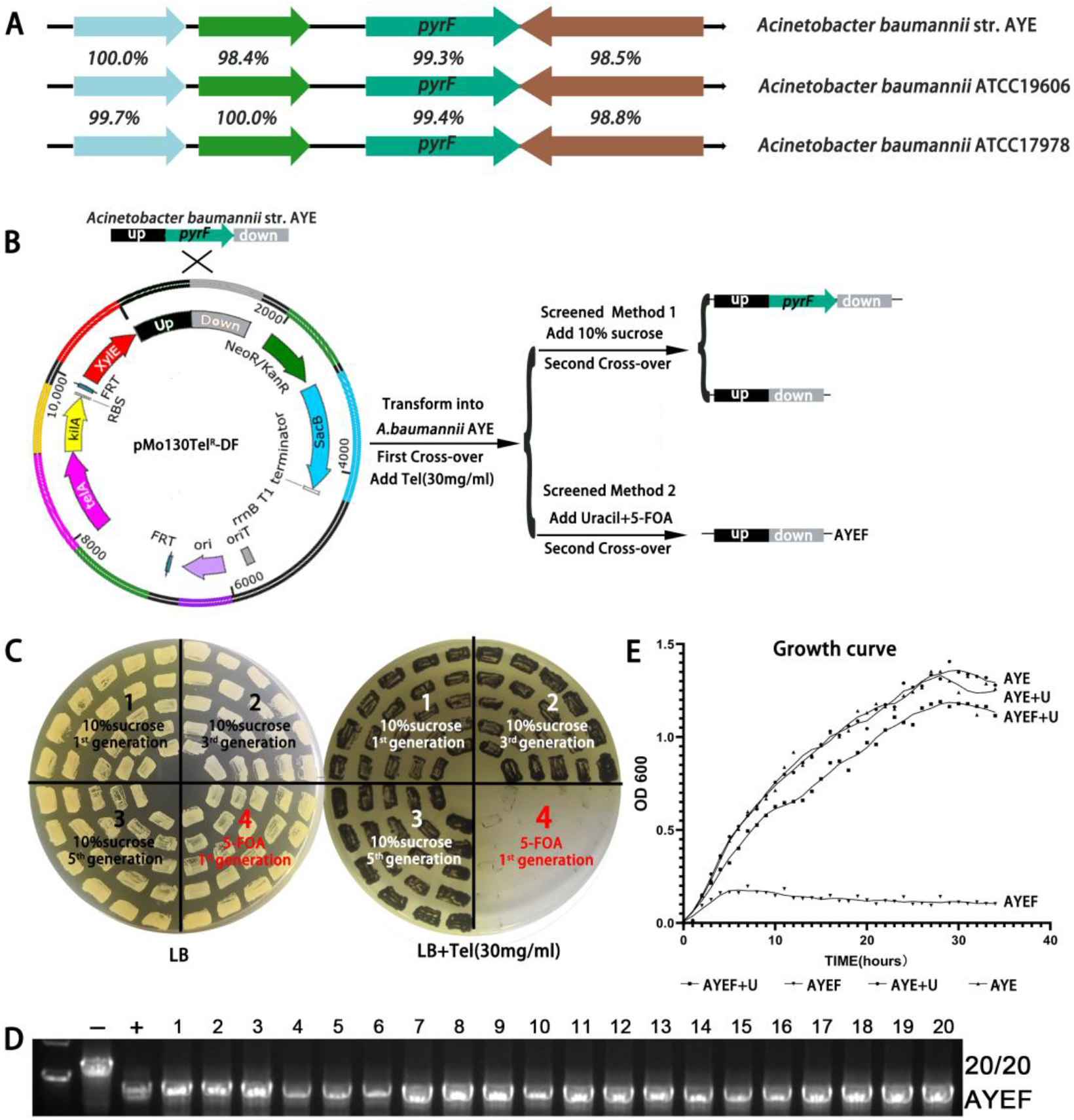
Clean marker-free *ΔpyrF* mutants were easily screened and constructed with 5-FOA in *A.baumannii* AYE. **A.** The *pyrF* gene and its flanking DNA sequences are highly conserved in *A.baumannii*. The *pyrF* and its flanking DNA sequences were analyzed by blastn (https://blast.ncbi.nlm.nih.gov/Blast.cgi) for most *A.baumannii*. The diagram only shows the results for three commonly used model strains. **B.** The schematic diagram of method for clean marker-free *ΔpyrF* mutants constructing in *A.baumannii* AYE. The plasmid pMo130Tel^R^-DF was generated by harboring the *pyrF* 500bp-flanking DNA fragments of AYE. The first-crossover mutants were screened by transforming and culturing in LB with Tel (30mg/L). The second-crossover mutants were screened by two different counter-selection methods (10% sucrose or 50mg/L 5-FOA). **C.** Photocopying plate method screened the second-crossover colonies. The Tel-susceptible colonies were the second-crossover colonies. Region 1, picked after 1^st^ generation with 10% sucrose added. Region 2, picked after 3^rd^ generation with 10% sucrose added. Region 3, picked after 5^th^ generation with 10% sucrose added. Region 4, picked after 1^st^ generation with 5-FOA (50mg/L) added. **D.** PCR screened for the second-crossover clones from Region 4 (5-FOA, 1.st generation). The second-crossover colonies were all *pyrF* gene deleted (20/20). **E.** Growth curve of the AYE and AYEF strains in different liquid medium (LB, LB+U)

#### The wild-type *A. baumannii* strains were all susceptible to 5-FOA and autonomously synthesized uracil

The uracil prototrophic and the sensibility to 5-FOA of *A. baumannii* are the preconditions for developing a *pyrF*-based selection and counter-selection system. A synthetic medium (M95) lacking uracil was first used for the uracil prototrophic, and the wild-type *A. baumannii* strains autonomously synthesized uracil and grew well in M95 medium (Fig. S1C, **M95**). But when we streak these strains on the LB plates with 5-FOA (50mg/L), they all susceptible to 5-FOA and could not grow (Fig. S1C, **LB+UF**). Thus, the *pyrF* gene was considered to be responsible for the uracil biosynthesis and 5-FOA sensitivity in *A.baumannii*. And the uracil auxotrophic and 5-FOA susceptible combined with *pyrF* gene could be broadly utilized as the selection (first-crossover) and counter-selection (second-crossover) for the genetic manipulation system in *A.baumannii*.

### Deletion the *pyrF* gene to generate the *ΔpyrF* mutants as the uracil auxotrophic host in *A.baumannii*

#### Clean marker-free *ΔpyrF* mutants were easily screened and constructed with 5-FOA

Firstly, we generated a *pyrF*-deleted mutant and named it AYEF by using the pMo130Tel^R^ system in *A. baumannii* AYE. A schematic drawing of the strategy used for deleting the *pyrF* gene in *A. baumannii* AYE is shown in Fig. 1B. The plasmid pMo130Tel^R^-DF carried the conserved 500bp-*pyrF*-flanking DNA fragments of AYE was generated and transformed into the strain *A. baumannii* AYE, which has integrated into the genome by the first-crossover homologous recombination with Tel (30mg/L) as the selection agent. The first-crossover colony (Tel^R^) was picked to generate the second-crossover colonies, with 5-FOA (50mg/L) as the counter-selection agent. 20 5-FOA resistant colonies were picked for photocopying plate streaking. These 5-FOA resistant colonies were all sensitive to Tel and second-crossover colonies (20/20, Fig. 1C, **Region 4**), which indicated that the *pyrF* gene and the plasmid derivatives had been excised from the genome by second-crossover homologous recombination. PCR amplification were performed for these second-crossover colonies and further confirmed the *pyrF* gene were all deleted (20/20, Fig. 1D). So it was easily to create a clean marker-free *ΔpyrF* mutant by this method. In comparison, we also used sucrose as the counter-selection agent, but after five subgenerations, the plasmid still difficult to be eliminated from the genome and the second-crossover colonies were hard to obtained (Fig. 1B, **Region 1, 2, 3**). What’s more, PCR amplification is also needed for screening the correct colonies.

As the 500bp-flanking DNA sequences of *pyrF* were highly conserved, we also generated the *ΔpyrF* mutants for *A. baumannii* strains 17978, 19606 and 11 clinical strains with the same plasmid pMo130Tel^R^-DF and the same method (Fig. S1A). And as results, the *pyrF* genes were all easily and successfully deleted (Fig. S1B). So the *ΔpyrF* mutant generated method may be also applicable to more *A. baumannii* strains.

#### *ΔpyrF* strains were resistant to 5-FOA and uracil auxotrophic

*A. baumannii* AYE and AYEF were cultured in LB liquid medium, the growth of AYEF was significantly inhibited, but it grew same as AYE with uracil added (Fig. 1E). For all the wild type and *ΔpyrF* mutants, LB plates contained uracil and 5-FOA (LB+UF), and M95 plates were used to detect the 5-FOA susceptible and uracil auxotrophic, with the LB+U plates and the M95+U plates as the control. The *ΔpyrF* mutants grew well and the wild-type strain could not grow in LB+UF plates (Fig. S1C), thus the *ΔpyrF* mutants were all resistant to 5-FOA. The wild-type strains grew well but the *ΔpyrF* mutants could not grow in M95 plates indicates that the *ΔpyrF* strains were all uracil auxotrophic. And with uracil added in M95 plates it also grew well. These results indicated that the deletion of *pyrF* gene limited the synthesis of uracil, and sequentially affected the growth, which could be complemented by adding back the exogenous uracil. The wild-type strains were susceptible to 5-FOA and the deletion of *pyrF* conferred resistance to 5-FOA, the wild-type strain synthesized uracil independently and the deletion of *pyrF* conferred uracil auxotrophic, which means that the *pyrF* gene mediated the uracil biosynthesis and 5-FOA sensitivity in *A.baumannii*.

### Generation the suicide and autonomously replication vector with the *pyrF* gene introduced

Besides the uracil auxotrophic host strain, plasmids used as the tools for gene manipulation also needed to be generated, with *pyrF* gene as a selectable or counter-selectable marker to be introduced into plasmids for function complement. A brief scheme of the plasmids construction is shown in Fig. 2.

**Figure 2.**
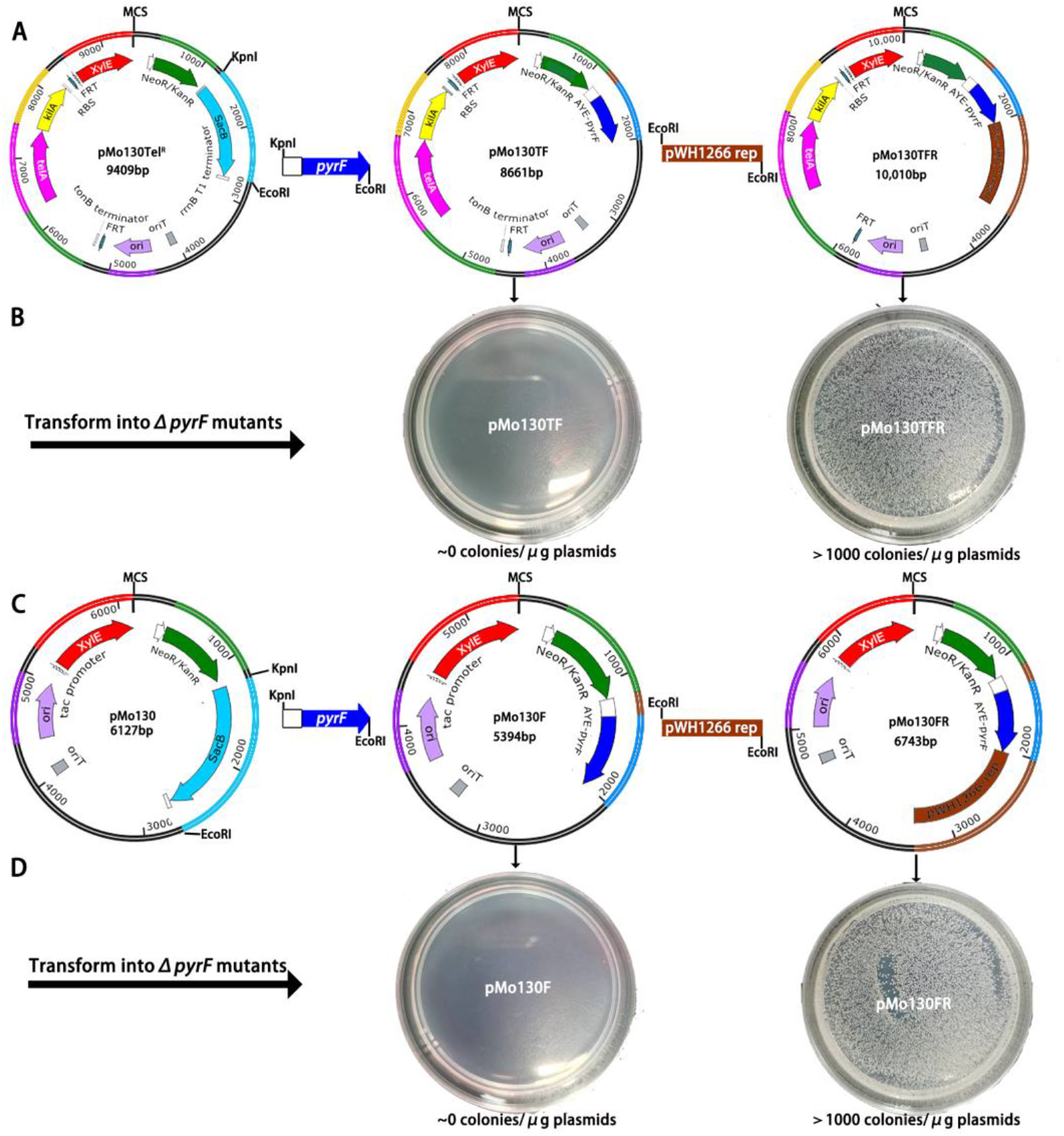
The *pyrF* gene as the selection marker or counter-selection marker was introduced into plasmids. **A.** The non-replicative plasmids pMo130TF and pMo130F were constructed with the AYE *pyrF* gene and its upstream 200bp promoter introduced into pMo-Tel^R^ and pMo130 by KpnI and EcoRI. **B.** pMo130TF and pMo130TFR were transformed into AYEF. Over 1000 colonies grew for the pMo130TFR, while none colonies grew for pMo130TF on M95 plates. **C.** The replicative plasmids pMo130TFR and pMo130FR were constructed with the pWH1266 *rep* introduced into pMo130TF and pMO130F by EcoRI. **D.** pMo130F and pMo130TFR were transformed into AYEF. Over 1000 colonies grew for the pMo130FR, while none colonies grew for pMo130F on M95 plates.

Non-replicative plasmids (suicide plasmids) were usually used for target sequence deletion, insertion, replacement or interruption. The pMo130TF and pMo130F suicide plasmids cannot replicate in *A.baumannii*, and was utilized to integrate into genome for creating single-crossover mutations. The *pyrF* gene (along with its 200bp-upstream promoter sequence) was introduced into pMo130Tel^R^ (Tel^R^ and Km^R^) or pMo130R (Km^R^) to replace the *sacB* gene to generate the pMo130TF and pMo130F (Fig. 2A). The *pyrF* gene could be used as the selectable marker for first-crossover in the uracil auxotrophic host strain when cultured in the M95 medium (uracil auxotrophic), and also used as the counter-selectable marker for second-crossover with 5-FOA added.

The replicative plasmid was usually used for gene expression or complement, which can autonomous replication without integrating to the chromosome. Two autonomous replication plasmid s pMo130TFR or pMo130FR (Fig. 2C) were generated by introduced the *rep* gene of pWH1266 into pMo130TF or pMo130F. The *pyrF* gene could also be used as the selectable marker for first-crossover in the uracil auxotrophic host strain when cultured in the M95 medium (uracil auxotrophic).

The pMo130F (5394bp) and pMo130FR (6743bp) were constructed for its much smaller size than pMo130TF (8661bp) and pMo130TFR (10010bp), which could be used to carry large DNA fragments for insertion or complement.

### Detection the *pyrF* gene function complements using the pMo130TFR and pMo130FR

As above designed, the pMo130TFR and pMo130FR were used to complement the *pyrF* gene function and could autonomous replicated. To verify these functions, the pMo130TFR (1μg) and pMo130FR (1μg) were transformed into AYEF or other *ΔpyrF* mutants, with pMo130TF (1μg) and pMo130F (1μg) as the negative control. As expected, there were over 1000 colonies for the pMo130TFR and pMo130FR, while there were none colonies for pMo130TF and pMo130F on M95 plates (Fig. 2B&D). This means pMo130TFR and pMo130FR could autonomously replicate without integrating to the chromosome, while pMo130TF cannot replicate without homologous sequence. And the pMo130TFR and pMo130FR trans formants grew well on the M95 plates also means that the *pyrF* gene in the plasmid could express and complement the function, which can be used as the selection with uracil auxotrophic and counter-selection marker with 5-FOA toxic effect.

### Verification of the *pyrF*-based selection and counter-selection system with gene manipulation of the CRISPR-Cas systemin *A. baumannii* AYEF

We had succeeded to construct the uracil auxotrophic host strain *A. baumannii* AYEF and the suicide vectors for homologous recombination. In order to demonstrate the usefulness and convenience of this *pyrF*-based selection and counter-selection system for gene manipulation, we used it to modify the *cas* genes and CRISPR sequence of I-F CRISPR-Cas system in *A. baumannii* AYEF. A schematic drawing of the strategy used to gene manipulation of these DNA sequences is shown in Fig. 3A&B. We generated pMo130TF-E*cas1*, pMo130TF-E*cas3*, pMo130TF-E*cade* and pMo130TF-ECRI to delete *cas1*, *cas3*, *cascade* and large part of CRISPR (spacer2 and the flanking downstream CRISPR sequence deleted). After electro transformation, the first-crossover colonies were obtained on the M95 plates. And then these colonies were cultured with uracil and 5-FOA added for counter-selection to screen the second-crossover colonies. After first-generation cultured with adding 5-FOA, the integrated plasmid of E*cas1* (20/20), E*cas3* (20/20), E*cade* (20/20), and ECRI (19/20) were efficiently removed from the genome (Fig. 3C, **Region 4**). PCR amplified were performed to further screen the correct gene deletion colonies for E*cas1* (8/20), E*cas3* (7/20), E*cade* (9/20), and ECRI (8/19), (Fig. 1D). As comparison, we also introduced these sequences into pMo130Tel^R^ system and used the sucrose as the counter-selection agent to gene manipulation of these above sequence. After five consecutive screenings with *sacB/*sucrose system, the plasmid of E*cas1* (11/20), E*cas3* (0/20), E*cade* (0/20), and ECRI (0/20) were still hard to be completely removed from the genome (Fig. 3C, **Region 1, 2, 3**). The plasmids integrated in genome were removal from genome efficiently for *pyrF*/5FOA system, while the *sacB*/sucrose system was difficult. We also insert a GST tag at the C-terminal of *cas3*, which was also efficiently and conveniently screened the correct colonies for the E*cas3-GST* (10/20), (Fig. 3D). So the *pyrF*-based selection and counter-selection system is an efficient and convenient choice for gene manipulation.

**Figure 3.**
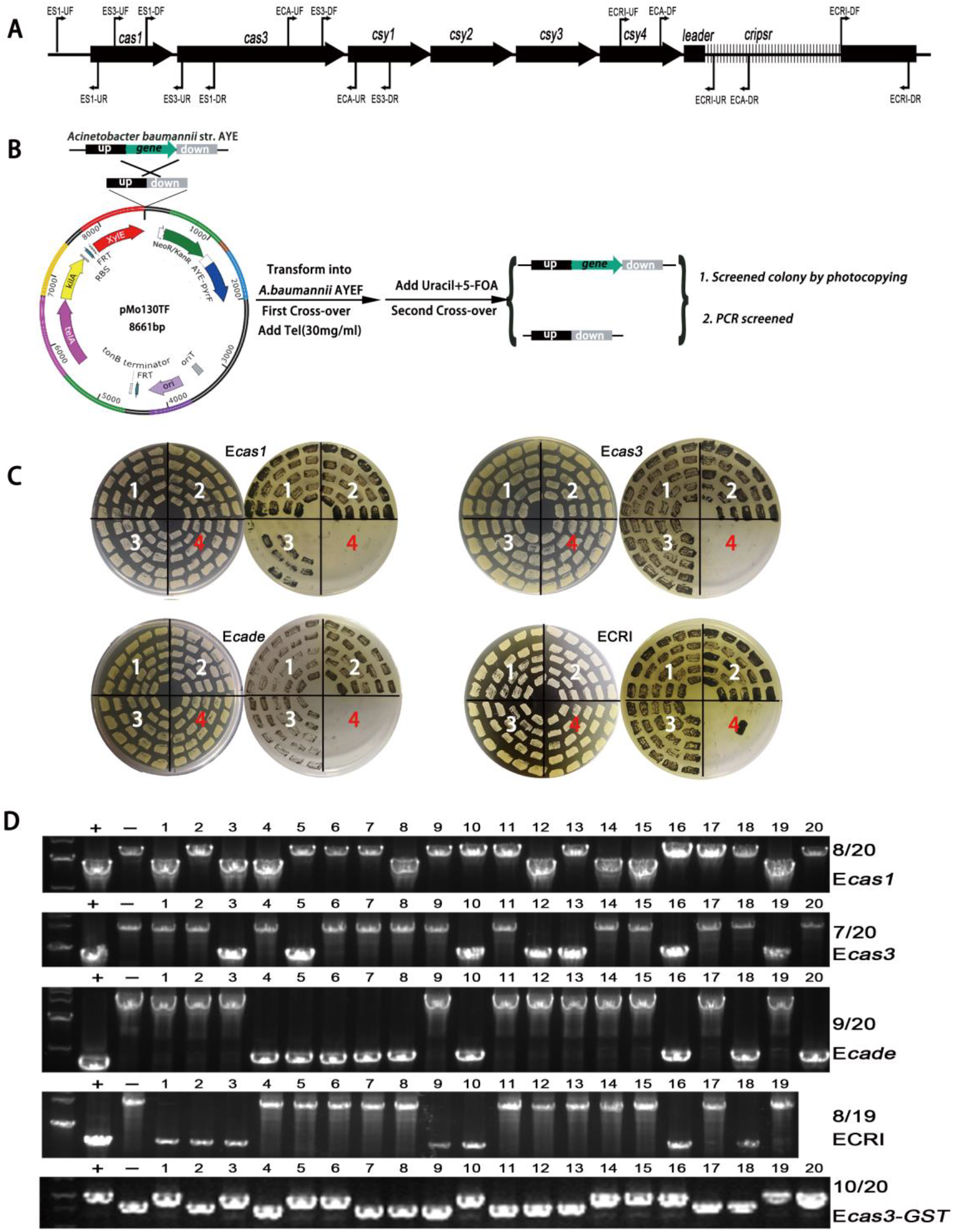
Verification of the *pyrF*-based selection and counter-selection system with gene manipulation of the CRISPR-Cas system in *A.baumannii* AYEF. **A.** The schematic diagram of the CRISPR-Cas system in *A.baumannii* AYEF. **B.** The schematic diagram for the target gene manipulation based on pMo130TF with *pyrF*/5-FOA as the counter-selection marker. **C.** Photocopying plate method screened the second-crossover colonies for E*cas1*, E*cas3*, E*cade*, and ECRI. The Tel-susceptible colonies were the second-crossover colonies. Region 1, picked after 1^st^ generation with 10% sucrose added. Region 2, picked after 3^rd^ generation with 10% sucrose added. Region 3, picked after 5^th^ generation with 10% sucrose added. Region 4, picked after 1^st^ generation with 5-FOA (50mg/L) added. **D.** PCR screened for the second-crossover clones from Region 4 (5-FOA, 1.st generation). E*cas1* (8/20), E*cas3* (7/20), E*cade* (9/20), ECRI (8/19) and E*cas3-GST* (10/20) were successfully constructed. -, wild type stains; +, plasmid; M, marker.

### Detection the CRISPR interference and adaptation in the AYEF and its mutants

Based on the high transformation efficiency of the replicative plasmid pMo130TFR, we generated the pMo130TFR-ECC-sp2 to verify the function of CRISPR-Cas system with the pMo130TFR as the control. A schematic drawing of the strategy used to detection is shown in Fig. 4A. If the CRISPR-Cas system works well, it would produce the s2-crRNA from the CRISPR, which can target the sequence ECC-spacer2 in the target plasmid pMo130TFR-ECC-sp2. Eventually, the CRISPR interference and adaptation would happen, when the necessary *cas* gene and DNA elements were completely existed. If the CRISPR interference works, the pMo130TFR-ECC-sp2 would be cut for targeted, and the number of transformed clones would decrease. So the CRISPR interference could be detected by counting the colonies and conducted comparative analysis between pMo130TFR and pMo130TFR-ECC-sp2 after transformation (Fig. 4B). The CRISPR adaptation could be detected for the colonies by PCR amplifying the sequence flanking the CRISPR leader side, and the expanded band (larger than the parental band) could be found for spacer insertion and then sequenced for confirming (Fig. 4C&D). As results, we found that lacking of *cas3*, *Cascade*, and *spacer2* were almost the same number of colonies between pMo130TFR and pMo130TFR-ECC-sp2, while AYEF or the lacking of *cas1* mutant was decreased obviously for pMo130TFR-ECC-sp2 (Fig. 4E). This means that *cas3*, *Cascade*, and *spacer2* were involved in interference. And the adaptation cannot happen when the lack of *cas1*, *cas3*, *cascade* and spacer2 (Fig. 4C&E), which means they are necessary, and the adaptation of I-F CRISPR in *A. baumannii* AYEF needs a priming process. The pMo130TFR-ECC-sp2 could generate the interference and adaptation also means the PAM-CC is an interference and adaptation PAM. As results, we found the I-F CRISPR-Cas system was functioned well; all Cas proteins excepted Cas1 were involved in interference, and almost all Cas proteins and a priming spacer were needed for adaptation.

**Figure 4.**
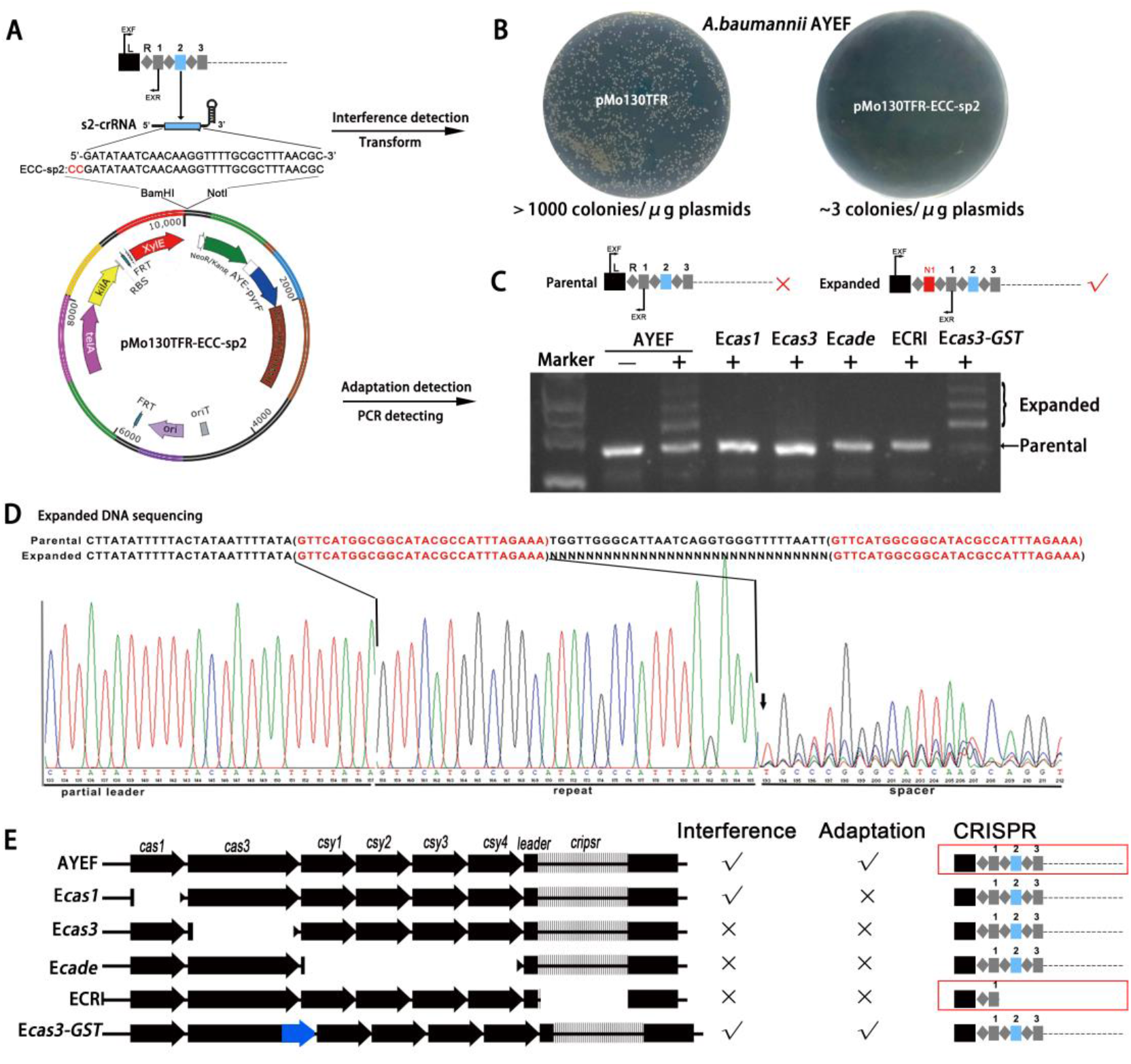
Detection of the CRISPR interference and adaptation in the AYEF and its mutants. **A.** The schematic diagram of the pMo130TFR-ECC-sp2 constructed and targeted. The s2-crRNA could be produced by the CRISPR of AYEF and then target the sequence ECC-spacer2 introduced into the pMo130TFR-ECC-sp2. **B.** Detection of CRISPR interference by transformation of the pMo130TFR-ECC-sp2, with pMo130TFR as the control. The number of transformed clones would be obviously decreased for pMo130TFR-ECC-sp2 (~3 colonies/μg plasmids) than pMo130TFR (> 1000 colonies/μg plasmids). **C.** Detection of the CRISPR adaptation by PCR amplify ing the sequence flanking the CRISPR leader side. The expanded bands (larger than parental band) were produced by spacer insertion colonies (adaptation). E*cas1*, E*cas3*, E*cade*, and ECRI could not detect the adaptation; AYEF and E*cas3-GST* could detect the adaptation -, pMo130TFR; +, pMo130TFR-ECC-sp2; M, marker. **D.** Chromatograph map that shows the sequencing result of the expanded DNA mixture in panel. **E.** The inductive results for CRISPR interference and adaptation in the AYEF and its mutants.

And that, for CRISPR adaptation when cultured the pMo130TFR-ECC-sp2 transformed colonies in the liquid M95 or in LB+Tel, the expanded PCR bands were gained more in M95 as the number of passages increases, while in LB+Tel the expanded PCR bands almost the same as the first subgeneration. This reveals that the uracil auxotrophic selection is better than the antibiotic selection marker in this situation (data not shown).

## Discussion

*A. baumannii* is a very urgent pathogenic microorganism, with great interest for public, due to the high mortality rates for nosocomial infection and high levels of extremely drug-resistant isolates. Convenient and quick genetic methods can help to understand the basic biology of *A. baumannii* infection and transmission, and also provide the molecular mechanism of gene function for these progresses. Recombineering method is a quick and efficient way to perform rapid genetic manipulation in vivo. For introduction of genetic material into *A. baumannii*, antibiotic selection markers were usually used. But for multi-drug-resistant *A. baumannii* strains, it was usually not so easy to find a suitable antibiotic marker and need to examine the antibiotic sensitivity and the minimal inhibitory concentration (MIC) of the target *A. baumannii* strain firstly [14, 17–19]. Thus, broadly selectable markers need to be developed. Hygromycin B was found early that have antimicrobial activity against a wide range of microorganisms and a broadly expressed hygromycin resistance (HmR) marker was developed for selection [16]. And later, Amin *et al*. demonstrated that tellurite as a non-antibiotic marker could be broadly used in most *A.baumannii* strains and developed a pMoTel^R^ vector[15]. Based on this vector, in our study, we also used tellurite as the selection marker to broadly generate the *pyrF*-deleted *A.baumannii* hosts.

For recombination-mediated genetic engineering, early methods always used an antibiotic marker to replace the gene of interest. But these methods not only bring antibiotic into genome, but also are not suited for multiple gene deletions. So most recently, the FLP-FRT system and counter-selection marker were developed to generate the marker-free mutants and conduct multiple gene editing. Tucker et al. developed a recombineering method with Rec_Ab_ as a rapid and efficient way for gene editing, which used an FLP/FRT recombinase system to engineer marker-less mutants [14]. But for this method, the plasmids (Rec_Ab_-carried and FLP-carried plasmids) needed to be transformed and naturally removed for twice, which is inconvenient and time consuming. Combined the Rec_Ab_ recombination system, the CRISPR-Cas9-Based genome editing was subsequently developed for *A. baumannii,* for which the positive clones could be easier to obtain and screen [18, 19]. But this method also needs to transform two plasmid pCasAb-apr and pSGAb-km for genome editing and then cure the plasmid with sucrose, and that it used two selectable antibiotic markers may also limit its applications in MDR and extensively drug-resistant. The other marker-less methods were mostly used the *sacB*/sucrose as the counter-selection, which need to be passaged daily for several days and also time consuming [15–17]. In this study, we demonstrated that the *pyrF*/5-FOA as the counter-selection system was useful and efficient, most of the time it just needs to be subcultured once with appropriate concentration of 5-FOA for counter-selection, while the *sacB*/sucrose system for plasmid eliminating from the genome was still difficult even after five subgenerations. Based on the *pyrF*-based system, we efficiently and easily modify the I-F CRISPR-Cas sequence in *A.baumannii* AYEF. And then we also easily detected the interference and adaptation of the CRISPR-Cas system with the replicative plasmid and its derivatives (pMo130TFR and pMo130TFR-ECC-sp2). As results, we found that this CRISPR-Cas system functioned well, all Cas proteins excepted Cas1 were involved in interference, and almost all Cas proteins and a priming spacer were needed for adaptation. Based on these results, we could also conduct further research on the molecular mechanisms of I-F CRISPR-Cas system in *A.baumannii.*

What’s more, the color of antibiotic sometimes interfere for optical detection, such as tellurite (black), tetracycline (yellow); and repeated use of antibiotics for stress screening may cause the emergence of resistant co lonies or not conductive to some experimental research. So this uracil auxotrophic system may be more suitable than antibiotic for studying some biological processes, such as the interference and adaptation of CRISPR-Cas system in this study. The replicative plasmid used the *pyrF* /uracil auxotrophic system as the selection, which ensured a more stable plasmid existence. The CRISPR adaptation could be detected more efficiently and conveniently for continuously and steadily acquiring the spacers from the plasmid.

For this genetic manipulation system, it was first needed to generate the *pyrF*-deletion mutant as the uracil auxotrophic host strain, which might to take some time. But utilizing pMo130Tel^R^-DF, tellurite and 5-FOA, the *pyrF*-deletion mutant could be broadly and easily generated as the uracil auxotrophic host strain for model and clinical *A.baumannii* strains. And the *pyrF*-carried plasmid (suicide plasmid or replicative plasmid) could also work well and broadly used as the gene manipulation tools with *pyrF* as the selection or counter-selection markers in these hosts. Above all, the *pyrF*-based system is an efficient and convenient system for broadly used in most *A. baumannii* strains to study the molecular mechanisms of drug-resistance, virulence and other gene functions.

## ACKNOWLEDGMENT

This work was supported by Grants from the National Natural Science Foundation of China (grant 31701078), the Research Fund of Chengdu Medical College (grant CYZ17-02), the Research Fund of Non-coding RNA and Drug Discovery Key Laboratory of Sichuan Province (grant FB19-03), College Students’ Innovative Entrepreneurial Training Plan Program (grant s201913705118)

## Supplementary materials

**Figure S1.**
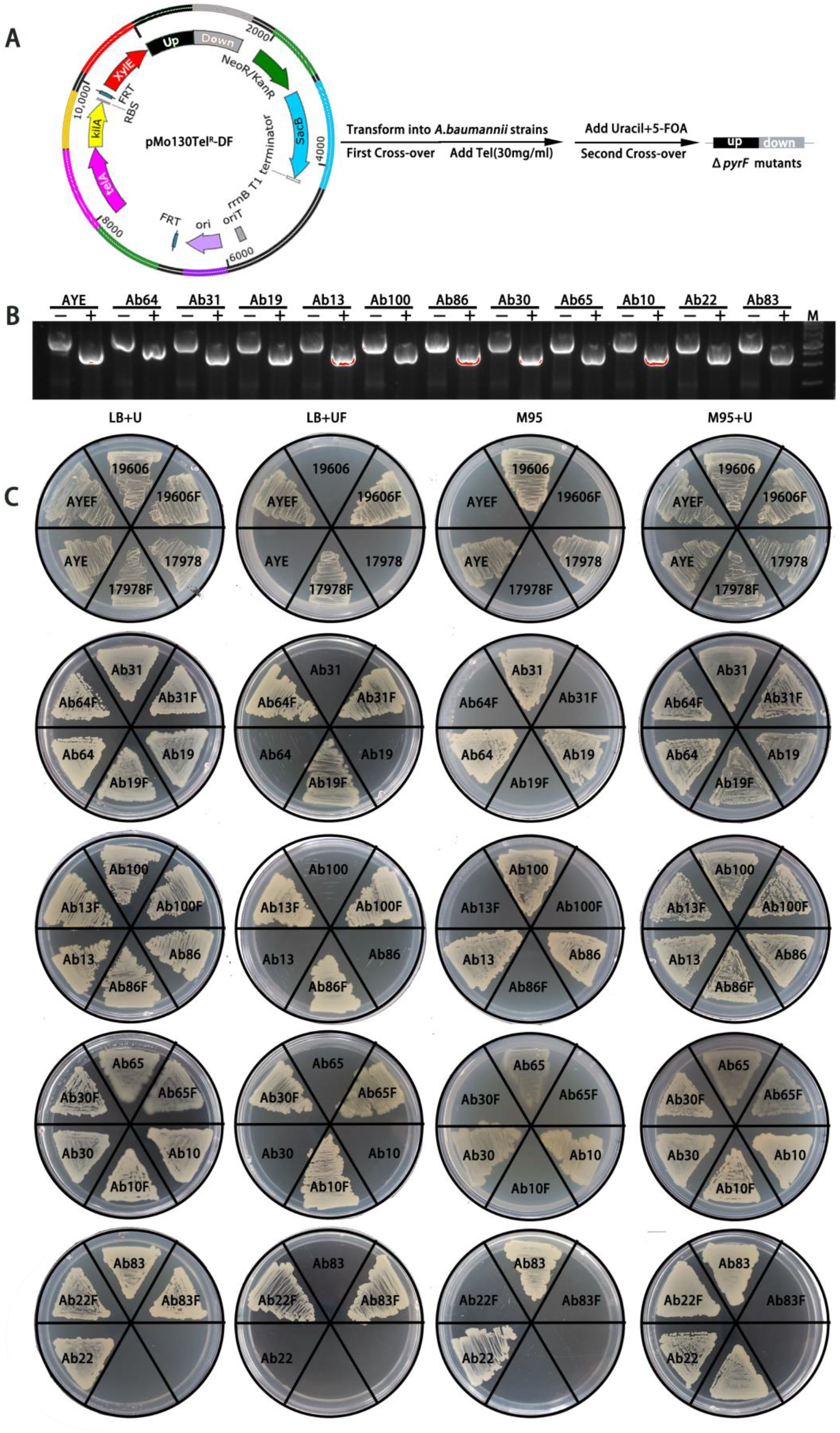
*ΔpyrF* mutants were easily screened and constructed with the pMo130Tel^R^-DF and 5-FOA in model and clinical *A.baumannii*. **A、** The schematic diagram of method for clean marker-free *ΔpyrF* mutants construction in model and clinical *A.baumannii*. Tel and 5-FOA were used as the selection and counter-selection agents to obtain the first-crossover and second-crossover colonies. **B、** PCR results shows the *pyrF* genes of 11 clinical strains were successfully deleted -, wild type stains; +, *pyrF*-deleted mutants; M, marker. **C、** The uracil prototrophic and the sensibility to 5-FOA for three model and 11 clinical strains *A. baumannii* and their *pyrF*-deleted mutants. LB+UF plates were detected the sensibility to 5-FOA, with LB+U plates as the control. M95 plates were detected the uracil auxotrophic, with M95+U plates as the control. Wild type strains: AYE, 19606., 17978., Ab64, Ab19, Ab13, Ab100, Ab86, Ab65, Ab30, Ab10, Ab22, and Ab83; *pyrF*-deleted mutants: Ab31F, Ab64F, Ab19F, Ab13F, Ab100F, Ab86F, Ab65F, Ab30F, Ab10F, Ab22F, and Ab83F.

**Table S1.**
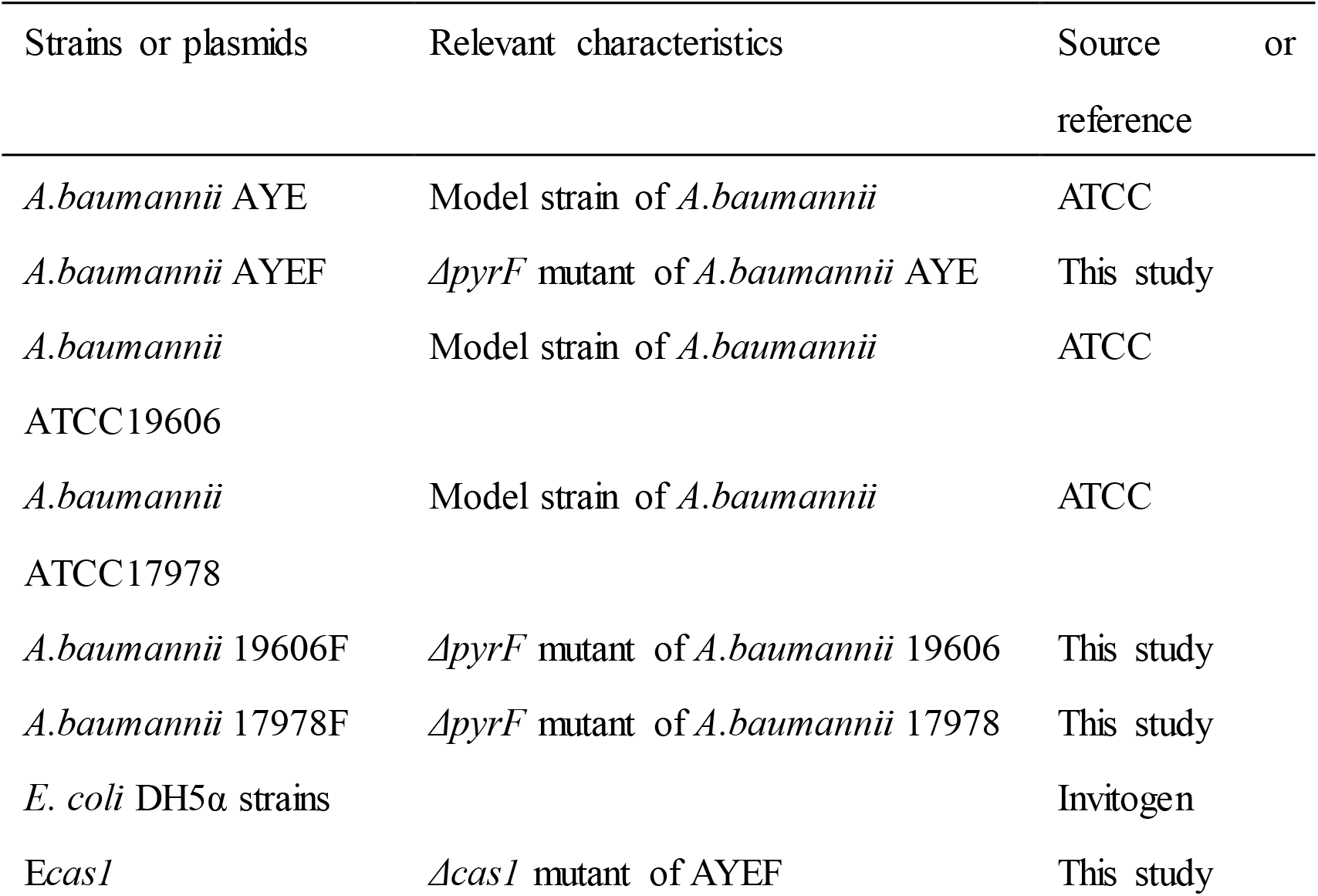
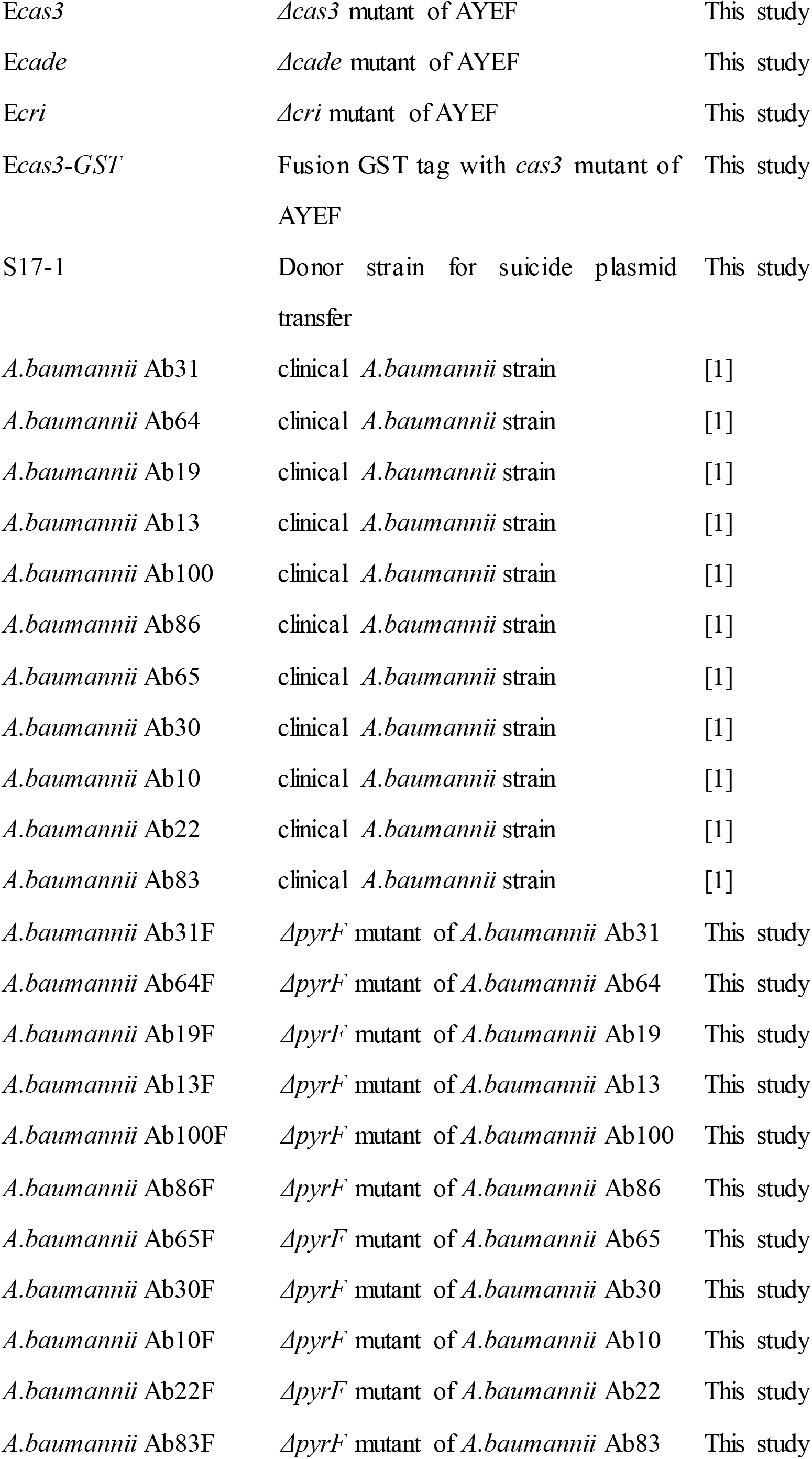
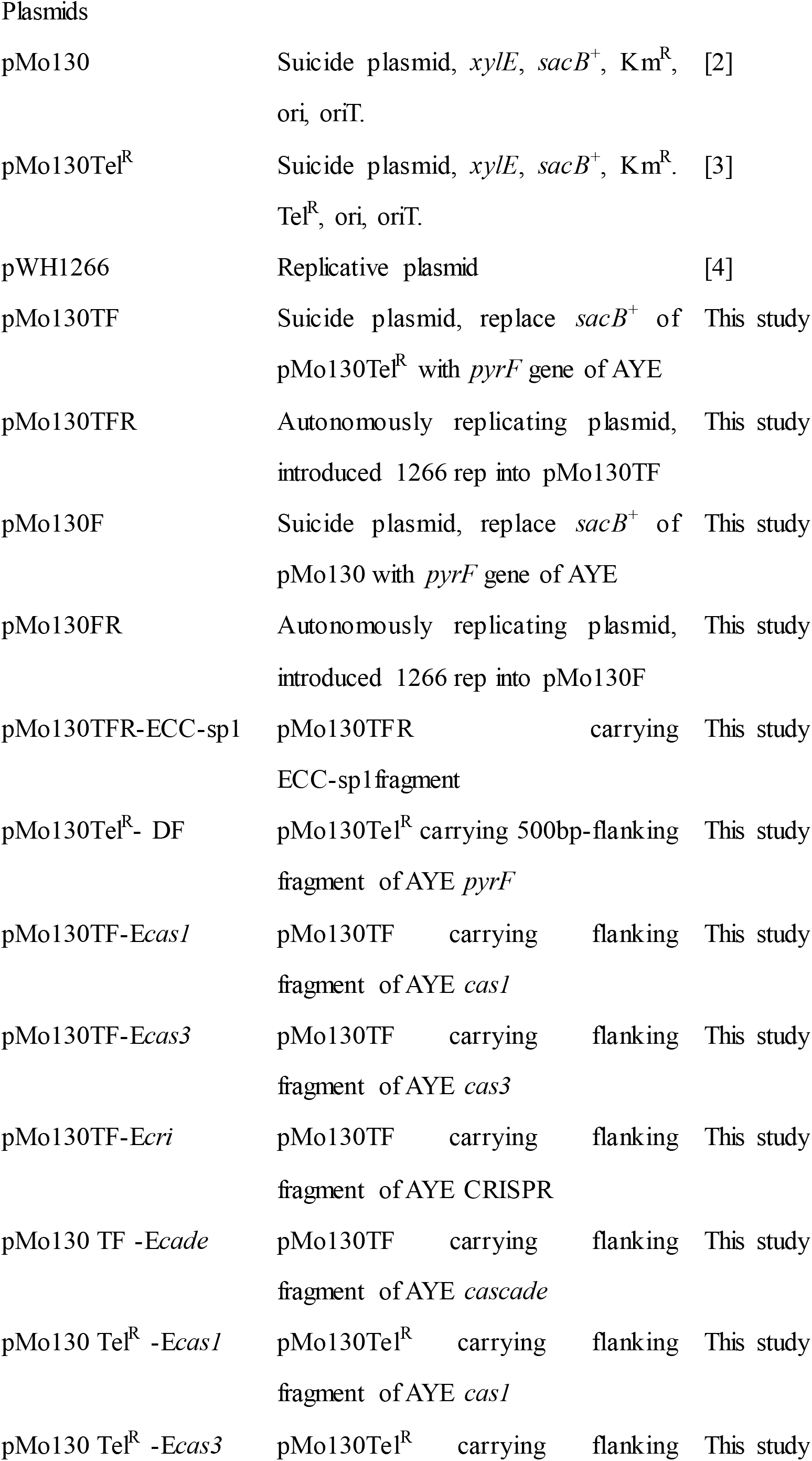
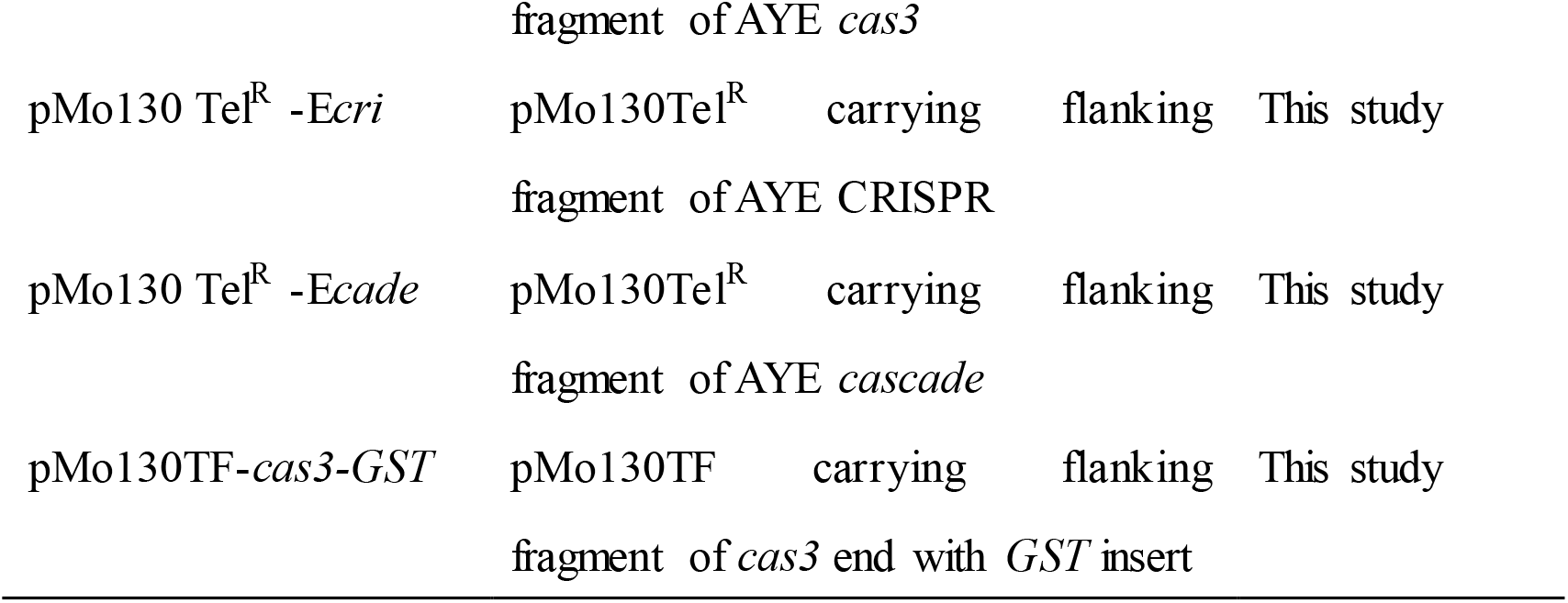
The strains and plasmids used in this study.

**Table S2.**
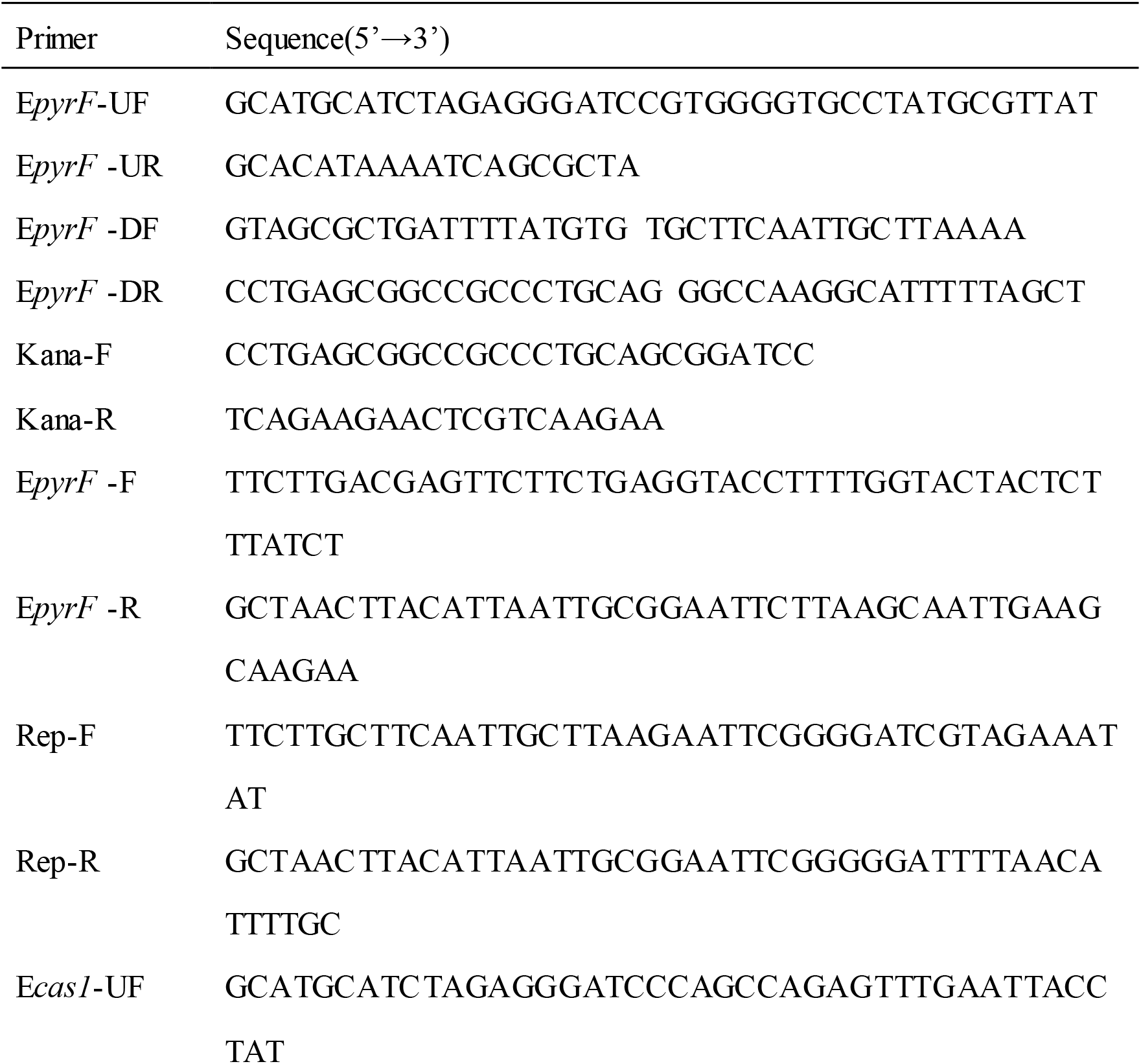
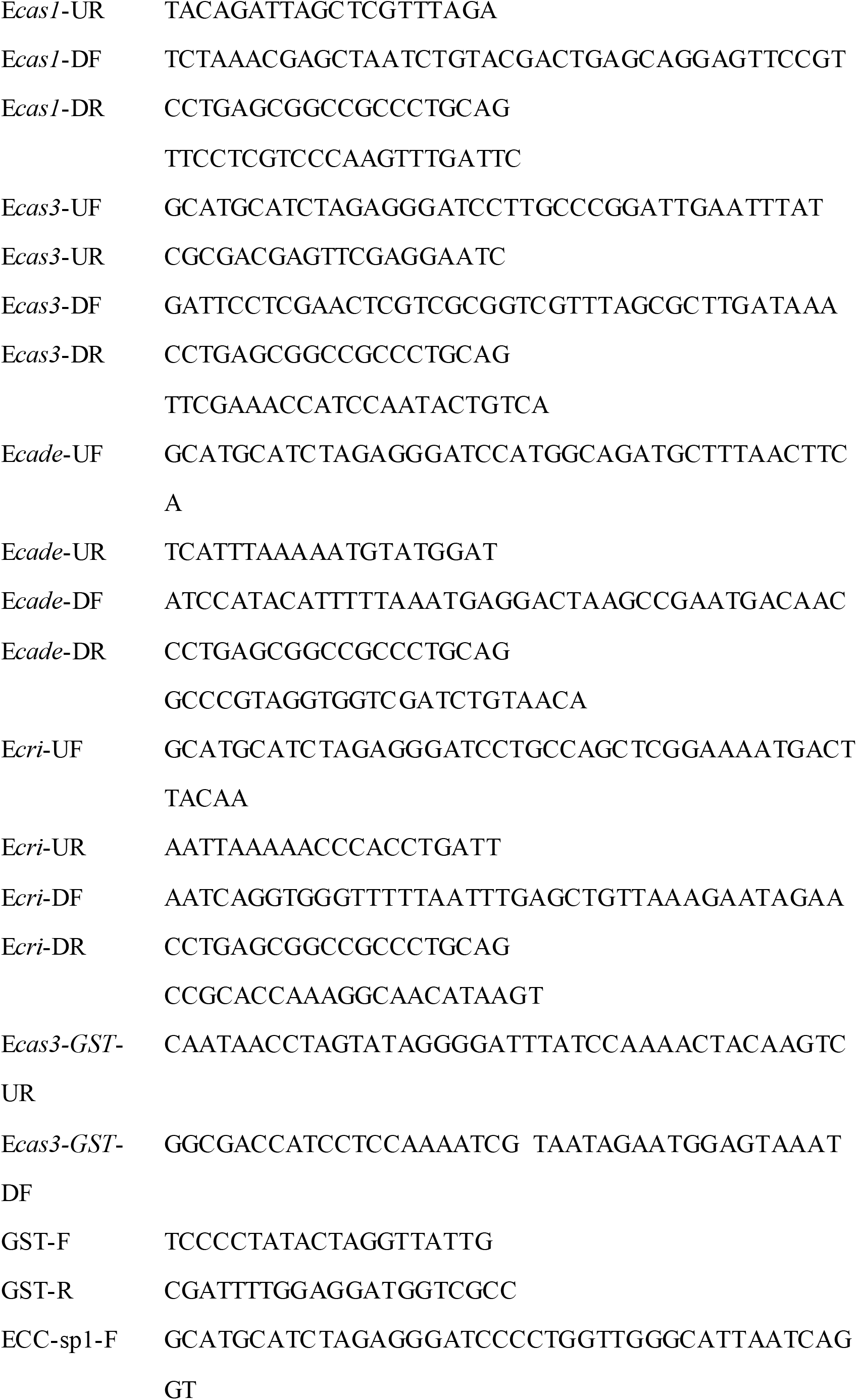
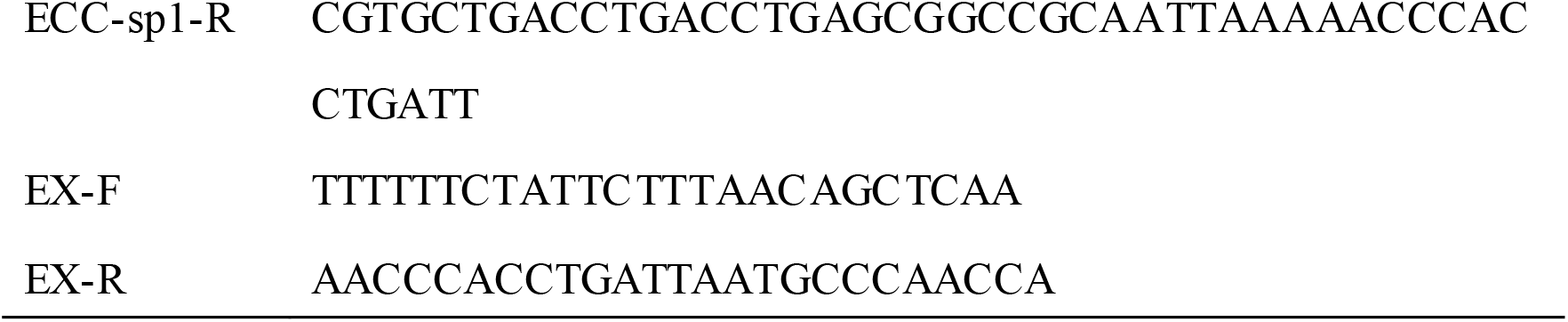
The primers used in this study.

